# Selective Inhibition of Facilitative Glucose Transporter 1 Improves Ultrafiltration Efficiency in Experimental Peritoneal Dialysis

**DOI:** 10.1101/2023.04.19.537457

**Authors:** Giedre Martus, Premkumar Siddhuraj, Jonas S. Erjefält, Martin Lindström, Karin Bergling, Carl M. Öberg

**Author notes:** **Correspondance:** Giedre Martus, Barngatan 2a, 221 85 LUND, Sweden, +46. **Funding** The study was funded by Lund University Medical Faculty Foundation grant YF 2020-YF0056, and The Inga-Britt and Arne Lundberg’s Research Foundation (all to C.M. Öberg).

## Abstract

Local and systemic side-effects of glucose remain major limitations of peritoneal dialysis (PD). Glucose transport during PD is thought to occur via inter-endothelial pathways, but recent data indicated that some glucose is transferred via facilitative glucose channels. Here we used BAY-876, a potent and highly selective blocker of facilitative glucose channel 1 (GLUT1), in an experimental rat model of PD using either 1.5% or 2.3% glucose fluid in a 1-h dwell. We also sought to elucidate whether diffusion of radiolabeled [^18^F]-deoxyglucose in the opposite direction (plasma → dialysate) is also lowered by selective/non-selective GLUT inhibition. Results show that selective GLUT1 inhibition markedly improved UF and enhanced the sodium dip, but no alterations in glucose transport or [^18^F]-deoxyglucose diffusion could be detected. Non-selective GLUT-inhibition using phloretin showed similar improvements on water and sodium transport, but also markedly decreased diffusion capacity for [^18^F]-deoxyglucose. We conclude that selective GLUT1 inhibition improved the UF efficiency in terms of mL of water removed per gram glucose absorbed by almost 70% for 1.5% glucose, implicating a role for GLUT1 in glucose mediated osmotic water transport in PD. Selective inhibitors of facilitative glucose transporter 1 may be promising agents to improve UF efficacy in patients treated with PD.

**Translational Statement:** Peritoneal dialysis (PD) is limited by systemic and local glucose toxicity. Here we used a highly selective inhibitor of facilitative glucose channel 1 (GLUT1) in a rat model of PD, and show marked, direct improvements in osmotic water removal (UF) per gram glucose absorbed. Inhibitors of GLUT1 may provide marked improvements in UF in patients on PD.

## Introduction

Glucose is added to peritoneal dialysis fluids to produce osmotic water removal across the peritoneal membrane. The presence of water channels in endothelial cells lining peritoneal capillary-and venular walls give rise to free-water transport accounting for about 40-50% of total UF ^1-3^, essentially doubling UF efficiency in terms of mL UF per gram glucose absorbed ^4^. Adverse effects from glucose exposure from peritoneal dialysis fluids, both systemic and local, remain important clinical problems, resulting in the development of metabolic complications ^5^, altering the structure and function of the peritoneal membrane ^6, 7^, and reducing the efficiency of the treatment -ultimately shortening time on therapy.

Glucose is impermeable across cell membranes, and is transported into cells via sodium-independent, facilitative glucose transporters (GLUTs) and sodium-glucose co-transporters (SGLT). Most cells express more than one kind of glucose transporter. Fourteen GLUT proteins have been described in the humans (GLUTs 1-14), and mediate the transmembrane movement of various monosaccharides and even myoinositol (GLUT13), urate (GLUT9), glucosamine (GLUTs 1/2/4), and vitamin C ^8^. Among the fourteen members in this family GLUT1-4 are the most extensively studied, and are believed to be most important for cellular glucose uptake. GLUTs have been detected in the peritoneal membrane ^9,10^. Schröppel *et al* demonstrated GLUT1 and GLUT3 expression in cultured human mesothelial cells and induced GLUT1 and 3 mRNA expression and glucose uptake by cells exposure to high glucose and cytokines ^11^. More recently, Schricker and colleagues found evidence of GLUT1 and GLUT3 expression in peritoneal membrane biopsies both by healthy controls, pre-dialytic patients, patients on PD and patients diagnosed with encapsulating peritoneal sclerosis, but analysis showed no significant differences in GLUT1 and 3 expressions between the different subgroups ^9^. Balzer *et al* showed in a mice model that GLUT1 and 3 was upregulated, but GLUT4 was downregulated in response to chronic glucose-based PD fluid exposure ^12^.

Our group recently demonstrated that phloretin, a non-selective GLUT blocker, reduced glucose absorption and improved ultrafiltration in a rat PD model ^13^. Due to the non-selective nature of phloretin, we could not answer the question via which GLUTs glucose absorption occurred, and whether GLUT-mediated glucose transport mainly occurs across or into cells. In order to cast light on these important questions, we investigated the effects of a potent (IC_50_ 2-3 nM), highly selective GLUT1 inhibitor BAY-876 using both 1.5% and 2.3% glucose (Figure 1). In a separate line of experiments, we studied the effects of GLUT-inhibition -using either BAY-876, phloretin and ritonavir (GLUT 1/4 blocker) -on the transport of ^18^F-deoxyglucose ([^18^F]-dg) in the opposite direction, from plasma to dialysate, during dialysis with 1.5% glucose fluids (Figure 1).

**Figure 1.**
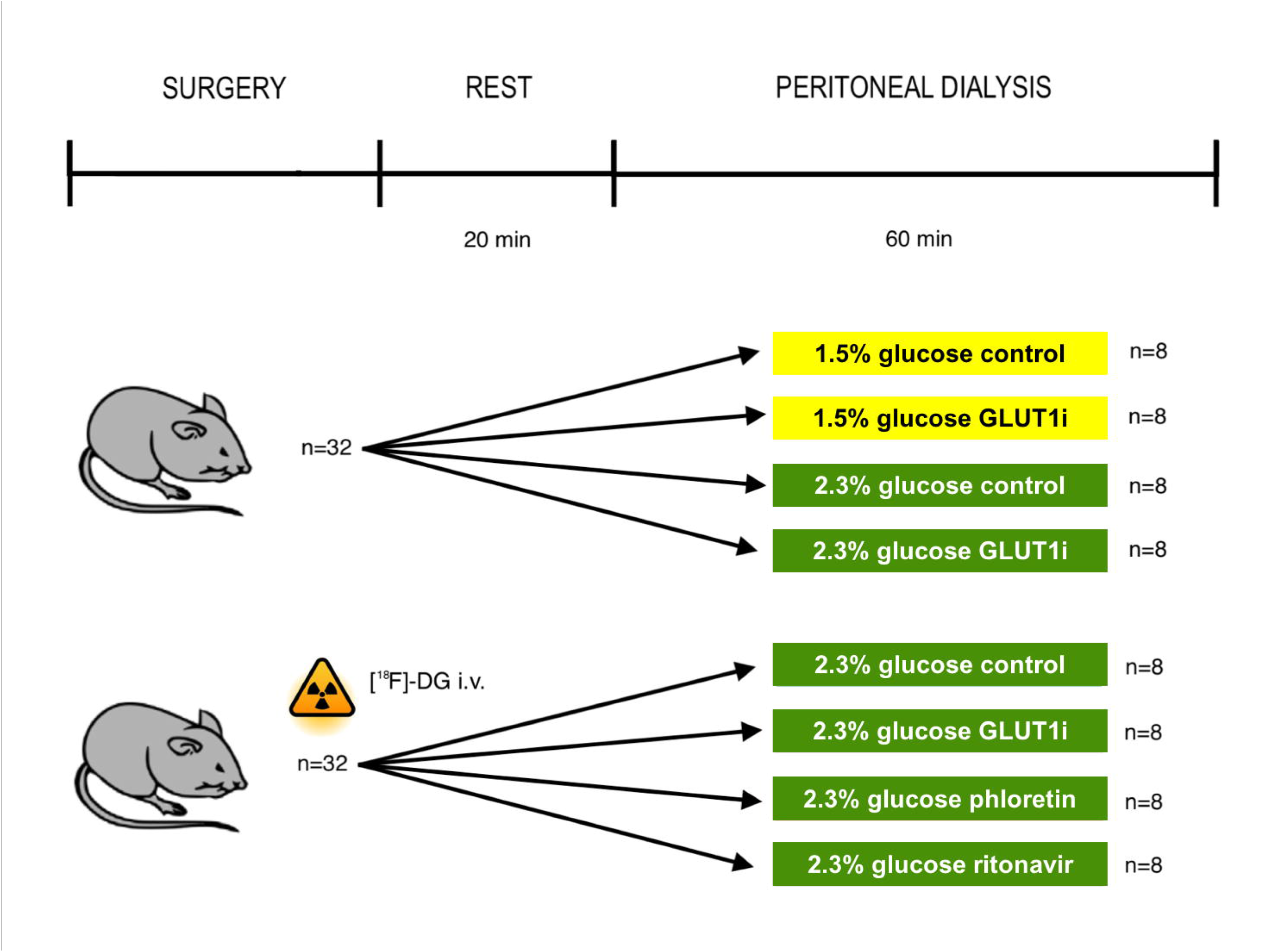
Schematic diagram of the experimental setup. Peritoneal dialysis was performed in anesthetized Sprague-Dawley rats using a fill volume of 20 mL with either 1.5% or 2.3% glucose fluid with or without selective GLUT1-blocker BAY-876, non-selective GLUT-blocker phloretin, and GLUT1/GLUT4-blocker ritonavir. Dialysate samples were obtained at baseline and after 30 min and 60 min dwell time. Routine blood samples were obtained before and after dialysis.

## METHODS

### Animals

Experiments were performed in male Sprague-Dawley rats having an average body weight of 304 g (215−348) with unrestricted access to water and food (Special Diets Services RM1(P) IRR.25 no. 801157). All animals were treated according to the guidelines of the National Institutes of Health for Care and Use of Laboratory animals. The Ethics Committee for Animal Research at Lund University approved of the experiments (Dnr 5.8.18-08386/2022). Exclusion criteria were signs of bleeding into the dialysate, premature death, signs of circulatory or respiratory failure, leakage of dialysis fluid or technical malfunction of the equipment. Animal care staff were unaware of the purpose of the experiments to ensure that all animals in the experiment are handled/treated in the same way. Reporting of results are in compliance with the ARRIVE guidelines. Each animal was gently placed in a covered glass container to which a continuous supply of 5% isoflurane in air (Isoban, Abbot Stockholm, Sweden) was connected. The animal was taken out of the container after being properly anesthetized. After that, 1.6-1.8% isoflurane in air was administered using a small mask to maintain the anesthesia. After tracheostomy, the animals were connected to a ventilator (Ugo Basile; Biological Research Apparatus, Comerio, Italy) and ventilated in a volume-controlled mode using a positive end-expiratory pressure of 4 cm H_2_O. Body temperature was kept between 37.1°C to 37.3°C via a feedback-controlled heating pad. End-tidal *p*CO_2_ was monitored continuously and kept between 4.8 and 5.5 kPa (Capstar-100, CWE, Ardmore, Pa). The left femoral artery was cannulated for monitoring of mean arterial pressure (MAP) and heart rate; and to obtain blood samples (95 μL) for measurement of glucose, creatinine, urea, electrolytes, hemoglobin, and hematocrit (I-STAT, Abbot, Abbot Park, Ill) before and after dialysis. The right femoral vein was cannulated and used for continuous saline infusion of 50 μL/min. The right internal jugular vein was cannulated for drug infusion. Access to the peritoneal cavity was established percutaneously via a multiholed silastic catheter (Venflon, BOC Ohmeda AB, Helsingborg, Sweden) (outer diameter 1.7 mm) secured to the skin using cyanoacrylate (Histoacryl, B. Braun Surgical, Rubi, Spain). After 60 minutes the dialysate was recovered from the peritoneal cavity, first by using a syringe, and thereafter carefully retrieving the rest of the fluid using pre-weighed gauze tissues. Wash-out was thereafter performed using 15 mL of PD-fluid with no tracer to retrieve residual amounts of tracer in the peritoneal cavity. As small amount of creatinine (∼0.3 mmol/L) was added to the dialysis fluid prior to infusion to allow dialysate concentrations to exceed the limit of detection of the iSTAT device. Hematocrit was determined by centrifugating thin capillary glass tubes. Peritoneal dialysis solutions were pre-warmed to 37°C before instillation. Animals were euthanized with an intravenous bolus injection of potassium chloride after the experiment. Prior to dialysate sampling, 1 mL was flushed back and forth several times to ensure a valid sample from the dialysate. Ultrafiltration rates (μL/min) was calculated from the total volume out – total volume in divided by 60 minutes. Net UF and solute mass balance (*e*.*g*., absorption) were determined as the mass (volume) out minus mass (volume) in. Plasma-to-dialysate clearances were calculated as the average solute removal rate (based on mass balance) divided by the mean plasma concentration (mmol/L). Dialysate clearances were calculated as the mean solute removal rate (based on mass balance) divided by the average dialysate concentration (mmol/L). Isovolumetric and isocratic diffusion capacities are calculated as described previously ^14^.

### Studies of GLUT1 inhibition

This study consisted of four groups of eight animals treated without or with BAY-876 (Merck, Darmstadt, Germany) 25 mg/L in 20 mL of either 1.5 % or 2.3% glucose peritoneal dialysis fluid (Balance, Fresenius, Bad Homburg, Germany). All animals were included for analysis. Dialysate samples for analysis of glucose concentration and electrolyte content (CHEM8) was obtained directly after instillation, at 30 min and after 60 min dwell time. Samples for radioactivity measurements (20 μL) were obtained from the dialysis fluid before infusion, and from the dialysate at 1, 10, 20, 30, 40, 50, and 60 min, and blood serum at 5, 15, 25, 35, 45, and 60 min and analyzed on a gamma counter (Wizard 1480; Wallac Oy, Turku, Finland) to determine ^51^Cr-EDTA and ^125^I-albumin activities. Urine was collected during dialysis and analyzed for glucose content (ISTAT) and radioactivity.^18^

### Studies of plasma-to-dialysate clearance of^18^ F-deoxyglucose

This study consisted of four groups of animals treated without intervention (SHAM; n=8) or with BAY-876 (25 mg/L; n=9), phloretin (50 mg/L; n=8) or ritonavir (60 mg/L; n=8) (all from Merck, Darmstadt, Germany) in 20 mL of 2.3 % glucose peritoneal dialysis fluid (Balance, Fresenius, Bad Homburg, Germany). One animal in the BAY-876 was excluded from analysis due to a malfunctioning infusion pump. On the day of the experiment, [^18^F]-dg was retrieved in the morning from the radiopharmaceutical preparation unit (part of the Department of Radiation Physics) at Skåne University Hospital in Lund in quantities of 200-500 MBq. A smaller amount of [^18^F]-dg was then added to saline to produce fluid for intravenous administration, used instead of maintenance fluids at a rate of 50 μL/min. Experiments were performed in collaboration with a radiation expert, and all staff handling [^18^F]-dg were equipped with dosimeters. Samples for radioactivity measurements (20 μL) were obtained from the dialysate at 1, 10, 20, 30, 40, 50, and 60 min, and blood plasma (obtained by centrifugation) at 5, 15, 25, 35, 45, and 60 min and analyzed on a gamma counter (Wizard 1480; Wallac Oy, Turku, Finland) to determine [^18^F]-dg activities. Since the half-life (T_half_) of [^18^F]-dg is only 109.734 min, measured activities were corrected for the decay at time *T* in minutes from the start of dialysis as follows

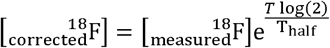

### GLUT1 Immunostaining

The surgically removed peritoneum tissue from rats was immediately transferred into buffered 4% formaldehyde (#02176; Histolab, Askim, Sweden) for 24 hours and subsequently tissue blocks were dehydrated in an automated dehydration machine (SAKURA-Tissue-Tek VIP® 6 AI; Sakura Finetek USA, Inc., Torrance, CA, USA). As part of the antigen retrieval process, slides (4 μm thickness) were baked at 60° C for 45 min and treated with a low pH target retrieval solution (#DM829; Dako, Glostrup, Denmark) in a DAKO PT Link HIER machine (PTlink 200; Dako, Glostrup, Denmark). The endogenous peroxidase enzyme was blocked with a peroxidase-blocking reagent (# DM821; DAKO, Glostrup, Denmark) for 10 minutes. Peroxidase-blocked slides were incubated with the primary antibody against GLUT1 (#SA0377; Thermo Fischer Scientific, Waltham, MA, USA) for 1 hour, followed by the secondary antibody (#DM822, Envision Flex, Dako, Glostrup, Denmark) for 30 minutes and finally substrate the HRP conjugated magenta dye (#DM857; DAKO, Glostrup, Denmark) for 10 minutes. For all the washing steps, readymade wash buffer (#DM831; Dako, Glostrup, Denmark) was used. After the immunostaining, the slides were counterstained with Mayer’s hematoxylin (#01820; Histolab, Askim, Sweden) and the coverslip (ECN 631-1574; VWR, Radnor, PA, USA) was mounted with Pertex (#00840; Histolab).

### Statistical methods

Data are shown as median (intraquartile range) unless otherwise indicated. Sample sizes were chosen to be similar to those in our previous experiments using GLUT-inhibitors ^13^. Significant differences were assessed using an omnibus test: non-parametric two-way ANOVA (ARTool) or a Kruskal-Wallis test as appropriate. P-values below 5% were considered significant for main effects, and below 10% for interactions. Corrections for multiple comparisons were performed when appropriate by using the Benjamini-Hochberg method. Calculations were performed using R for mac version 4.1.1.

## RESULTS

### Selective GLUT1 inhibition markedly increases ultrafiltration and 30 min sodium dip

UF rates were markedly improved in GLUT1 inhibitor treated animals both in the 1.5% and 2.3% glucose group, being 57 µl/min (IQR 52 to 59) compared to control group 35 µl/min (IQR 31 to 40) in the 1.5% glucose groups and 83 µl/min (IQR 80 to 86) compared to control 65 µL/min (IQR 61 to 69) in 2.3% glucose groups (Table 1) (Figure 2 a). Parallel to the improvements in UF, the 30 min sodium dip was increased in both groups (Figure 2 b).

**Table 1.**
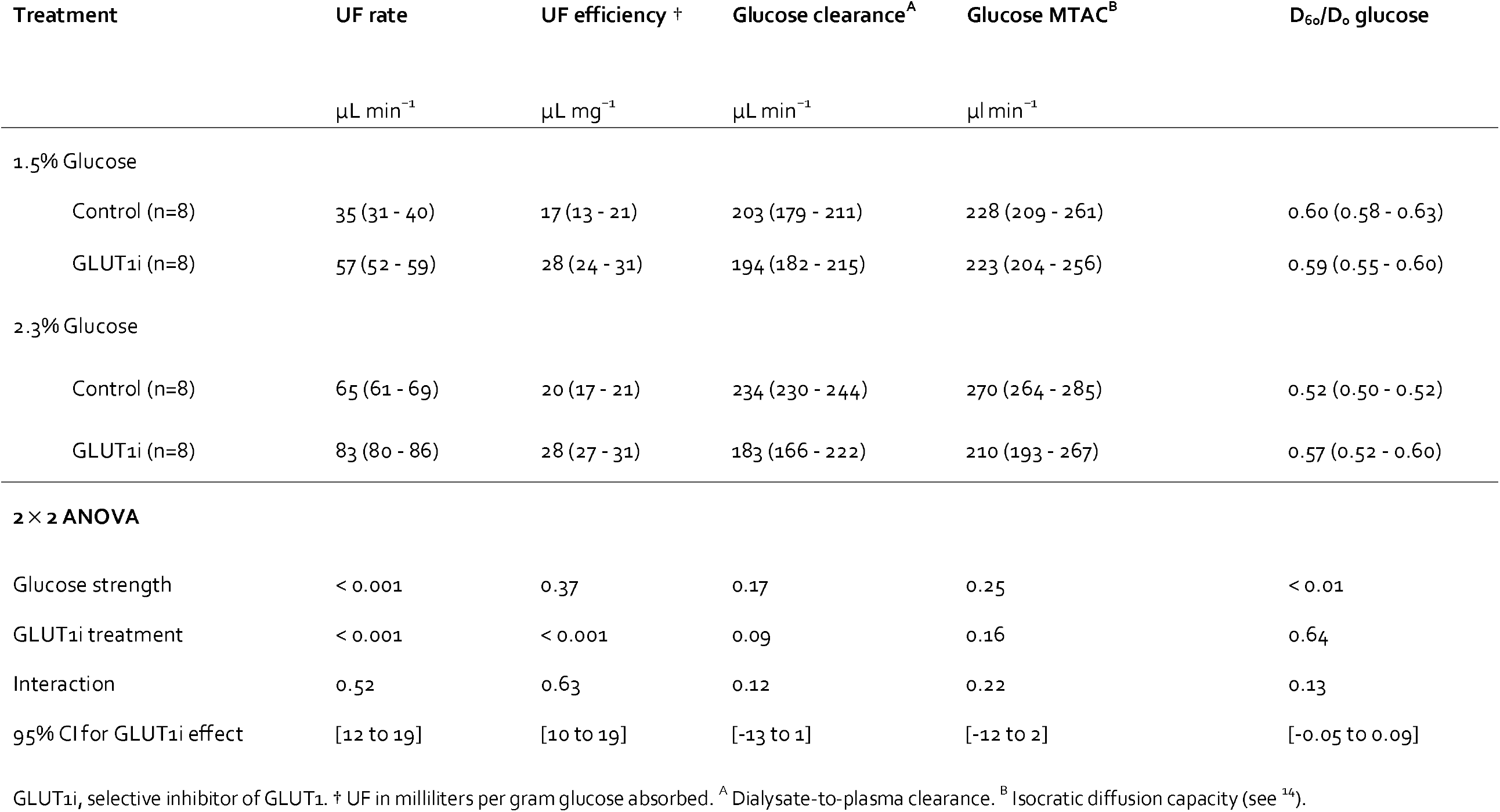
Effects of selective GLUT1 inhibition using BAY-876

| Treatment | UF rate | UF efficiency † | Glucose clearance<sup>A</sup> | Glucose MTAC<sup>B</sup> | D<sub>60</sub>/D<sub>0</sub> glucose |
| --- | --- | --- | --- | --- | --- |
|  | μL min<sup>-1</sup> | μL mg<sup>-1</sup> | μL min<sup>-1</sup> | μl min<sup>-1</sup> |  |
| 1.5% Glucose |  |  |  |  |  |
| Control (n=8) | 35 (31 - 40) | 17 (13 - 21) | 203 (179 - 211) | 228 (209 - 261) | 0.60 (0.58 - 0.63) |
| GLUT1i (n=8) | 57 (52 - 59) | 28 (24 - 31) | 194 (182 - 215) | 223 (204 - 256) | 0.59 (0.55 - 0.60) |
| 2.3% Glucose |  |  |  |  |  |
| Control (n=8) | 65 (61 - 69) | 20 (17 - 21) | 234 (230 - 244) | 270 (264 - 285) | 0.52 (0.50 - 0.52) |
| GLUT1i (n=8) | 83 (80 - 86) | 28 (27 - 31) | 183 (166 - 222) | 210 (193 - 267) | 0.57 (0.52 - 0.60) |
| **2 × 2 ANOVA** |  |  |  |  |  |
| Glucose strength | < 0.001 | 0.37 | 0.17 | 0.25 | < 0.01 |
| GLUT1i treatment | < 0.001 | < 0.001 | 0.09 | 0.16 | 0.64 |
| Interaction | 0.52 | 0.63 | 0.12 | 0.22 | 0.13 |
| 95% CI for GLUT1i effect | [12 to 19] | [10 to 19] | [-13 to 1] | [-12 to 2] | [-0.05 to 0.09] |

GLUT1i, selective inhibitor of GLUT1. † UF in milliliters per gram glucose absorbed. <sup>A</sup> Dialysate-to-plasma clearance. <sup>B</sup> Isocratic diffusion capacity (see <sup>14</sup>).

**Figure 2.**
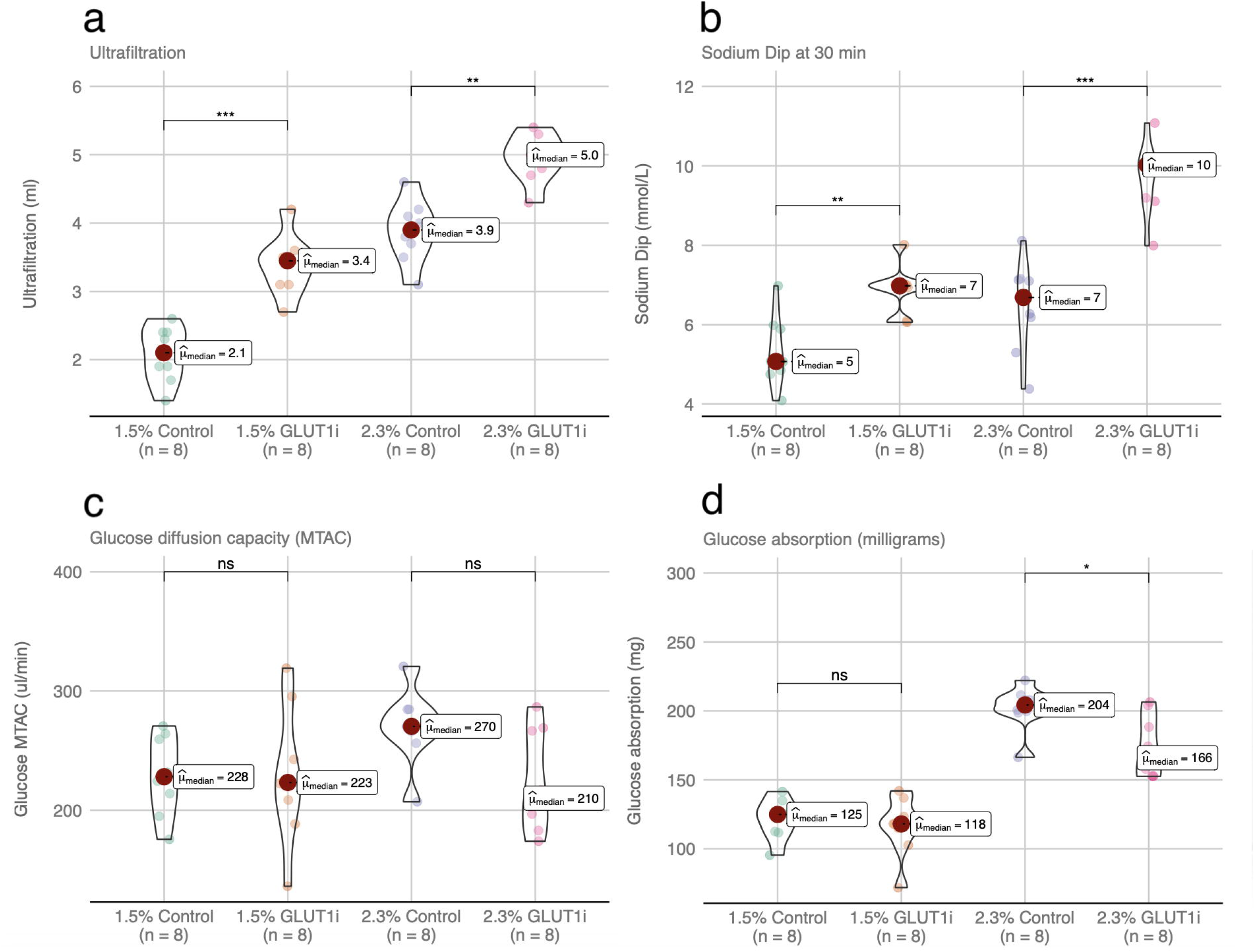
Selective GLUT1 inhibition markedly enhanced osmotic water removal and sodium dip. (a): Ultrafiltration (Volume out-Volume in), (b): Sodium dip at 30 min, (c): glucose isocratic diffusion capacity (see text), (c): Parietal peritoneum (Hematoxylin-Eosin stain), and (d): Glucose absorption (Glucose out – Glucose in) for the different groups.

### No significant effect of selective GLUT1 inhibition on glucose diffusion capacity

Isocratic diffusion capacities (MTAC) calculated for the 60 min glucose were not significantly different from between animals treated with GLUT1 inhibitor BAY-876 and controls (Figure 2 c) (Table 1). Similarly, isovolumetric (Henderson-Nolph) diffusion capacities appeared unaffected by GLUT1 inhibition, being 252 µl/min (IQR 231 to 286) compared to control 228 µl/min (IQR 227 to 246) in the 1.5% glucose groups and 266 µl/min (IQR 244 to 310) compared to 296 µl/min (IQR 286 to 309) in control animals for the 2.3% glucose groups. Glucose absorption was significant for GLUT1i treatment main effect (F_1,28_ = 5.99, p=0.02), and post-hoc tests showed significant differences only between the 2.3% glucose groups (P=0.024) (Figure 2 d). Fractional dialysate (D/D_0_) glucose concentration at 60 min was not significantly different in BAY-876-treated animals to controls both in the 1.5% and 2.3% glucose group, being 0.57 (IQR 0.54 to 0.62) compared to control group 0.57 (IQR 0.55 to 0.59) in the 1.5% glucose group and 0.51 (IQR 0.47 to 0.57) compared to control 0.47 (IQR 0.45 to 0.49) in 2.3% glucose group, *P* = 0.297 with an observed 95% CI -3.36 to 10.6.

### GLUT1 is mainly localized to the endothelium of peritoneal blood vessel walls

Immunohistochemical staining of GLUT1 in biopsy specimens from the peritoneum (Figure 3 a) using specific antibodies revealed diffuse positivity for GLUT1 in the endothelium of vessel walls, while the mostly denudated surface didn’t show any reaction in mesothelial cells, nor any reaction in myocytes or noticeable reaction in fibroblasts (Figure 3 b).

**Figure 3.**
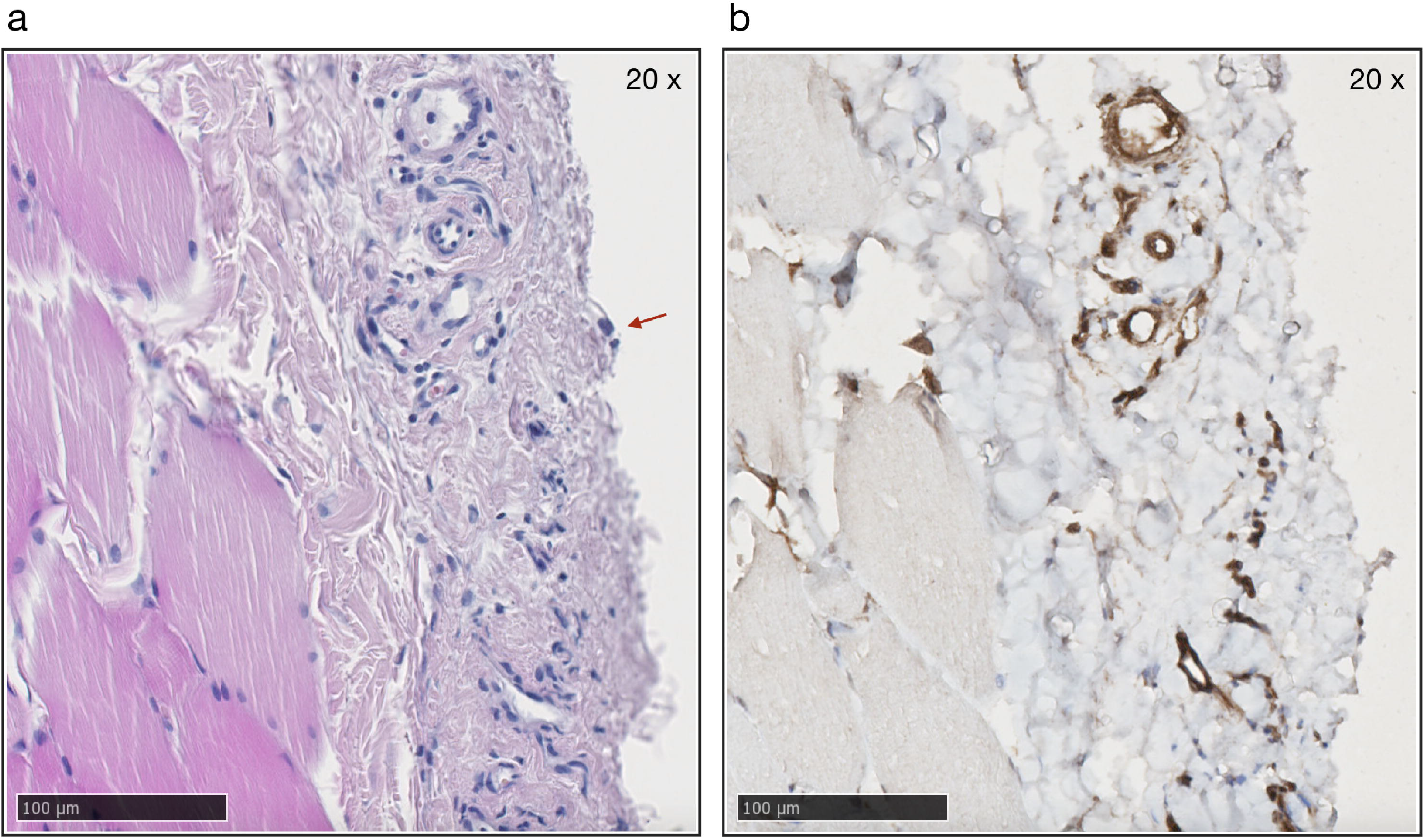
GLUT1 is localized to the endothelium of peritoneal blood vessel walls. (a): Parietal peritoneum (hematoxylin and eosin stain x 20) with several small vessels. Note single mesothelial cell (arrow). (b) Immune-staining of parietal peritoneum for GLUT1 (brown). Positive reaction in endothelium of small vessels shown in picture. Note absence of reaction elsewhere.

### Non-selective GLUT inhibition using phloretin reduced diffusion of radio-labelled [^18^F]-deoxyglucose whereas selective GLUT1 inhibition had no effect

The radionuclide [^18^F]-deoxyglucose ([^18^F]-dg) is widely applied for positron emission tomography imaging. The released positron is immediately annihilated upon collision with an electron, producing gamma radiation. Being a gamma radiator, [^18^F]-dg has the advantage of providing a lower error of measurement than conventional glucose measurements. Since error of measurement (combined with a low *N*) may be a possible explanation for the lack of a significant effect on glucose MTAC in the previous experiments, we next chose to study [^18^F]-dg transport in the same model. We also added groups receiving non-selective GLUT-inhibitor phloretin, as well as ritonavir (a GLUT1/GLUT4-blocker). After entering the cell via the same pathways as glucose, [^18^F]-dg is converted by hexokinase to [^18^F]-deoxyglucose-6-phosphate which is essentially trapped in the cell. Thus, one may expect that inhibiting cellular uptake in the peritoneal membrane would improve diffusion of [^18^F]-dg. In contrast, during a 60 min dwell utilizing 2.3% glucose, we noted marked *decrements* in [^18^F]-dg diffusion capacity during non-selective inhibition of GLUT channels using phloretin, but apparently not during selective GLUT1 inhibition (Figure 4) (Table 2). Such a decrease in [^18^F]-dg diffusion capacity indicate the possibility that a large part of [^18^F]-dg may be transported in a trans-cellular fashion across GLUTs other than GLUT1 in these experiments. Glucose diffusion capacities and [^18^F]-dg diffusion capacities were closely correlated (R2=0.49), but glucose MTACs were not significantly different between groups (Figure 4). No effects were observed regarding glucose plasma concentration or any other parameter before or after dialysis (Supplemental Table 1).

**Table 2.** Effects of non-selective GLUT-inhibition and selective GLUT1 inhibition on [^18^F]-deoxyglucose transport

| Treatment | UF rate | UF efficiency † | [<sup>18</sup>F]-dg MTAC<sup>A</sup> | Glucose MTAC<sup>A</sup> | Sodium dip |
| --- | --- | --- | --- | --- | --- |
|  | μL min<sup>-1</sup> | μL mg<sup>-1</sup> | μL min<sup>-1</sup> | μl min<sup>-1</sup> | mmol L<sup>-1</sup> |
| 2.3% Glucose |  |  |  |  |  |
| Control (n=8) | 75 (69 - 78) | 24 (22 - 30) | 206 (183 - 241) | 196 (173 - 239) | 8 (8 - 8) |
| GLUT1i (n=8) | 88 (86 - 92) | 32 (30 - 35) | 197 (179 - 205) | 197 (174 - 224) | 10 (9 - 11) |
| Phloretin (n=8) | 93 (88 - 98) | 36 (32 - 43) | 141 (134 - 163) | 154 (140 - 205) | 12 (11 - 12) |
| Ritonavir (n=8) | 88 (87 - 90) | 36 (33 - 39) | 164 (157 - 174) | 171 (138 - 183) | 10 (10 - 11) |
| **Kruskal-Wallis, p-value** | < 0.001 | 0.004 | 0.005 | 0.24 | < 0.001 |
| **Post-hoc analysis** |  |  |  |  |  |
| GLUT1i vs. control | 0.004 | 0.03 | 0.38 | - | 0.005 |
| Phloretin vs. control | 0.002 | 0.003 | 0.009 | - | < 0.001 |
| Ritonavir vs. control | 0.002 | 0.003 | 0.08 | - | 0.002 |

GLUT1i, selective inhibitor of GLUT1. † UF in milliliters per gram glucose absorbed. <sup>A</sup> Isocratic diffusion capacity (see <sup>14</sup>).

**Figure 4.**
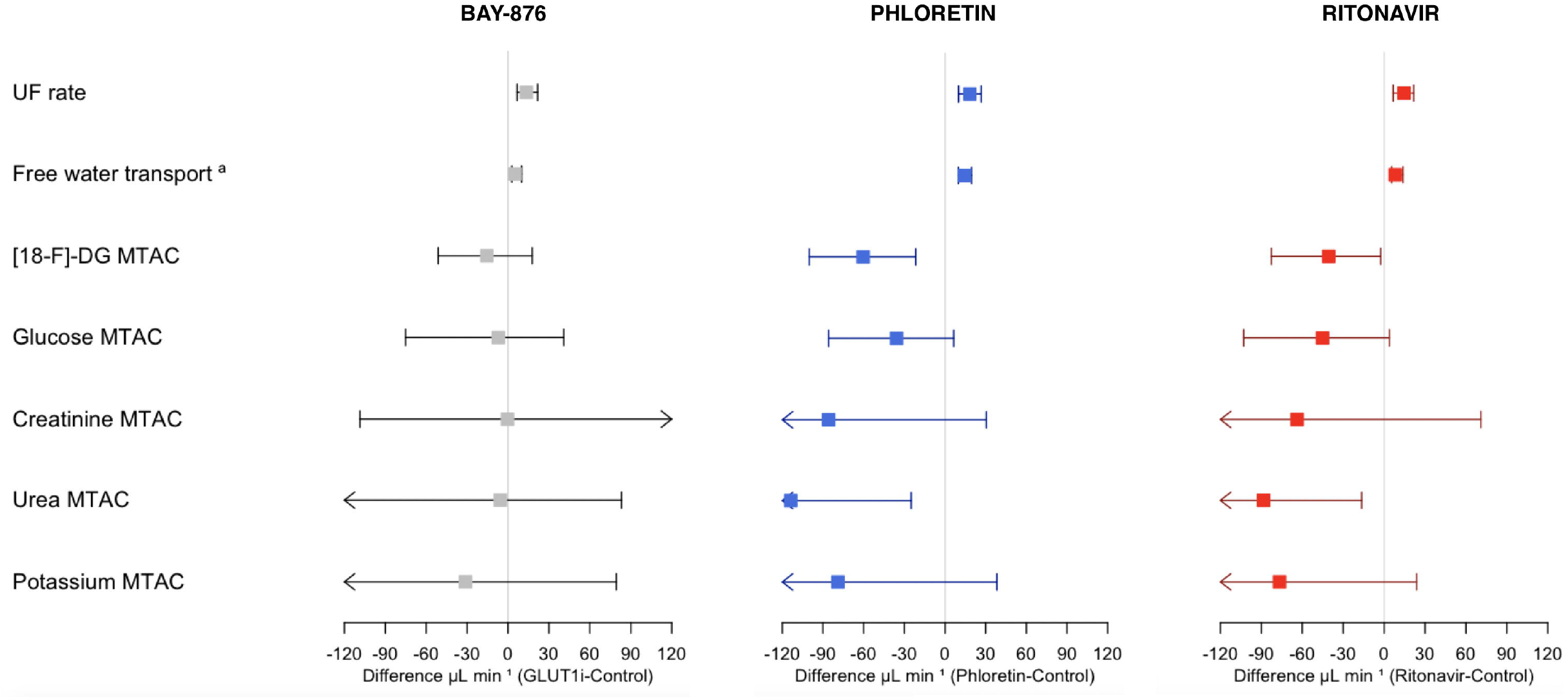
Treatment effects for selective GLUT1-inhibitor BAY-876, non-selective GLUT-inhibitor phloretin and non-specific GLUT1/4-blocker ritonavir. 95% CIs of the drug effects on UF rate, free-water transport, and isocratic diffusion capacities (MTAC) for [^18^F]-deoxyglucose, glucose, creatinine, urea, and potassium. Square boxes represent medians of the difference (drug-control). ^a^ Approximated as UFR-Sodium clearance.

### Three-pore model analysis

The parallel improvements in UF rate and sodium dip during selective GLUT1 inhibition without significant alterations in glucose diffusion capacity suggests that the ultrafiltration coefficient (LpS; water flux per mmHg of effective transmembrane pressure) was improved across the peritoneal membrane. This hypothesis is supported by three-pore model (Supplemental Table 2) transport parameter estimation, where the ultrafiltration coefficient was improved in both glucose strengths (Table 3). Again, glucose diffusion capacity was not significantly altered in the presence of selective GLUT1 inhibition (Table 3). Analysis of 2.3% glucose [^18^F]-dg data (Table 4) showed that phloretin effectively decreased diffusion capacity of [^18^F]-dg diffusion capacity, while there were no apparent effects on [^18^F]-dg diffusion capacity in animals treated with BAY-876 nor Ritonavir. Also, there were no significant effects on ultrafiltration coefficients and glucose MTACs in [^18^F]-dg animals treated with phloretin, ritonavir or BAY-876 (Table 4).

**Table 3.** Three-pore model analysis of selective GLUT1 inhibition effects on water and glucose transport

| Group | UF coefficient †<br>$\mu\text{L min}^{-1}\text{mmHg}^{-1}$ | Glucose MTAC<sup>A</sup><br>60 min $\mu\text{L min}^{-1}$ |
| --- | --- | --- |
| 1.5% Glucose |  |  |
| Control (n=8) | 1.2 (1.0 - 1.3) | 216 (182 - 236) |
| GLUT1i (n=8) | 2.1 (2.0 - 2.2) | 207 (143 - 224) |
| 2.3% Glucose |  |  |
| Control (n=8) | 1.2 (1.2 - 1.3) | 272 (259 - 277) |
| GLUT1i (n=8) | 1.6 (1.4 - 1.6) | 223 (184 - 262) |
| **2 x 2 ANOVA** |  |  |
| Glucose strength *p*-value | <0.001 | 0.066 |
| GLUT1i treatment *p*-value | 0.01 | 0.073 |
| Interaction *p*-value | <0.001 | 0.54 |
| 95% CI for GLUT1i effect | [1 to 2] | [-12.9 to 0.4] |

GLUT1i, selective inhibitor of GLUT1. † Also denoted LpS or Kf. <sup>A</sup> Three-pore model diffusion capacity

**Table 4.** Three-pore model effects of non-selective GLUT-inhibition and selective GLUT1 inhibition on [^18^F]-deoxyglucose transport

| Treatment group | UF coefficient †<br>μL min<sup>-1</sup> mmHg<sup>-1</sup> | Glucose MTAC<sup>A</sup><br>60 min μL min<sup>-1</sup> | [<sup>18</sup>F]-dg MTAC<sup>A</sup><br>60 min μL min<sup>-1</sup> |
| --- | --- | --- | --- |
| 2.3% Glucose |  |  |  |
| Control (n=8) | 1.4 (1.3 - 1.5) | 183 (156 - 209) | 198 (168 - 229) |
| GLUT1i (n=8) | 1.6 (1.5 - 1.7) | 192 (172 - 201) | 185 (164 - 199) |
| Phloretin (n=8) | 1.5 (1.5 - 1.7) | 185 (162 - 215) | 135 (122 - 160) |
| Ritonavir (n=8) | 1.6 (1.4 - 1.6) | 166 (157 - 178) | 155 (144 - 168) |
| **Kruskal-Wallis *p*-value** | 0.08 | 0.65 | 0.008 |
| **Post-hoc analysis** |  |  |  |
| GLUT1i vs. control *p*-value | - | - | 0.382 |
| Phloretin vs. control *p*-value | - | - | 0.009 |
| Ritonavir vs. control *p*-value | - | - | 0.057 |

† Also denoted LpS or Kf. <sup>A</sup> Three-pore model diffusion capacity

## DISCUSSION

The main finding in this study is that selective GLUT1 channel inhibitor BAY-876 increases ultrafiltration by ∼50% using 1.5% glucose – improving the UF efficiency in terms of mL of water removed per gram glucose absorbed by almost 70%. These results support the recent findings by our group using non-selective GLUT blocker phloretin ^13^, and also our previous findings using phlorizin ^15^ (which is converted to phloretin). Since the magnitude of the effects of GLUT1 inhibition alone approached that of non-selective GLUT-inhibition, it appears that a substantial part of the effects on UF in ^13, 15^ are due to inhibition of GLUT1. Additional mechanistic studies using [^18^F]-dg failed to detect differences in transport of both [^18^F]-dg and glucose during selective GLUT1-inhibition, indicating that the improvements in UF were apparently uncoupled from changes glucose transport across the membrane. This is despite the fact that the ultrafiltration rates along with marked increments in sodium dip indicate that the osmotic gradient was improved during selective GLUT1 inhibition, leading both to increased UF and free-water transport. In contrast, diffusion capacity of [^18^F]-dg was reduced during non-selective GLUT inhibition using phloretin. Taken together these results suggest that the mechanisms for glucose transport as well as osmotic water transport elicited by glucose across the peritoneal membrane are more complex than previously thought.

Actual transcellular transfer of glucose across endothelial cells in peripheral capillaries has as yet not been formally demonstrated ^16^. The present data nevertheless suggest the direct transport of glucose across cells, since we find a marked decrease in transport from plasma to dialysate during non-selective GLUT inhibition. However, in order for [^18^F]-dg to be transported *across* a cell, it must evade enzymatic phosphorylation by hexokinase, since it would otherwise be trapped inside the cell. It is possible that hexokinase is saturated during the rather extreme glucose concentrations during peritoneal dialysis. However, one early study by Betz *et al* have shown that not all [^18^F]-dg is converted by hexokinase, and 30-60% remains unphosphorylated ^17^. The next question is via which transporters such transcellular passage of [^18^F]-dg would occur. The lack of effect when blocking GLUT1 suggest that pathway is not involved, especially since the dialysate concentration of BAY-876 was ∼0.05 mM being several orders of magnitude larger than the IC_50_ of ∼2.5 nM. It is well-recognized that GLUT1 is the primary transporter for glucose across the blood-brain-barrier, whereas apical SGLT1 and basolateral GLUT2 is responsible for transcellular transport of glucose in the small intestine enterocytes ^16^. Given that peritoneal endothelia are not as tight as the blood-brain-barrier, the role of transcellular transport across peripheral capillary walls is likely less important.

When it comes to UF and glucose diffusion capacity our previous studies suggest that SGLT-channels appear to have no direct role under the experimental conditions in this study. Our previous experiments using a highly selective SGLT2-blocker empagliflozin ^14^ or a potent selective SGLT1-blocker mizagliflozin ^13^ both failed to show any effects on peritoneal water and glucose transport. Nevertheless, SGLT2-inhibition may have beneficial effects in the PD population ^18^, and chronic experimental models have demonstrated reduced peritoneal fibrosis in animals treated with SGLT2-inhibition ^12^. The limitations in this study are mostly related to the low number of animals in each group and the acute nature of the experiments. Thus, long-term effects cannot be evaluated from these data. Also, the glucose measurements have an error (VC) of about 6% which likely contributes to variance between groups. Lastly, the experiments using [^18^F]-dg compared three interventions (BAY-876, ritonavir and phloretin) with a single control (SHAM) group which may have decreased statistic power to detect significant differences.

We conclude that data so far indicate a role for GLUT transporters in peritoneal glucose transport – but which transporters other than GLUT1 are involved? Further experimental studies using more selective blockers may shed more light on these questions. The present study supports a partial role for GLUT1, and possibly GLUT4 since results using GLUT1/4 blocker ritonavir appeared to be more effective than GLUT1-inhibition alone. However, ritonavir may have other effects unrelated to GLUTs, which makes the interpretation of these results difficult. Given the existence of several medications with an inhibitory action on GLUTs, and the fact that the expression in peritoneal tissues of these transporters appear to be increased during PD, clinical trials to assess the therapeutic effects of glucose transport inhibitors in PD patients are warranted as the next step. Our recent pre-clinical study ^13^ indicates that circa 20-30 % of glucose may be transported via GLUTs. This is in contrast with clinical results in a previous study from our group showing that creatinine and glucose diffusion capacities in PD-patients were highly correlated ^19^. In contrast, one would expect a systematic bias between the MTAC for creatinine and glucose if a constant portion of glucose was transported via another pathway across glucose transporters. Nevertheless, it is possible or even likely that the expression of glucose transporters in a clinical cohort is highly variable which perhaps could explain the rather high variability in glucose MTAC ^19^. In the same study it is evident that water transport (i.e. osmotic conductance) and solute transport (creatinine MTAC) appear to be uncoupled in PD-patients. Intriguingly, the present study also supports such an uncoupling since it appears that water permeability was markedly improved during GLUT1 inhibition with little or no effect on solute transport. In conclusion our data supports a role for GLUT1 in the regulation of osmotic water transport during PD.

## Data sharing statement

All original data reported in this article have been deposited in Dryad (doi:10.5061/dryad.2547d7wwc).

## Supplemental material

Supplemental material for this article is available online. Supplemental Table S1, Supplemental Table S2, Supplemental Table S3, Supplemental Table S4.

## Acknowledgements

The authors are very grateful for the excellent experimental work by Helén Axelberg. We are also grateful for all helpful advice from radiation expert Hanna Holstein, and the excellent people at the department of Medical Radiation Physics at Lund University.

