## Supplemental material for "Selective Inhibition of Facilitative Glucose Transporter 1 Improves Ultrafiltration Efficiency in Experimental Peritoneal Dialysis"

Supplemental Material File Listing

Supplemental Table S1

Supplemental Table S2

**Supplemental Table 1.** Systemic effects of GLUT inhibitiors (Δ=after-before dialysis)

| **Treatment** | **Δ Plasma glucose**  mmol L^−1^ | **Δ Plasma creatinine**  μmol L^−1^ | **Δ Plasma sodium**  mmol L^−1^ | **Δ Plasma potassium**  mmol L^−1^ | **Δ Plasma TCO_2_**  mmol L^−1^ | **Δ Plasma iCa**  mmol L^−1^ |
| --- | --- | --- | --- | --- | --- | --- |
| 1.5% Glucose |  |  |  |  |  |  |
| Control (n=8) | 1.2 (0.5 - 2.0) | 8 (6 - 13) | 2.0 (1.0 - 2.0) | -0.1 (-0.3 - 0.0) | -1.0 (-1.2 to -1.0) | 0.06 (0.02 - 0.07) |
| GLUT1i (n=8) | 1.4 (-0.3 - 2.4) | 6 (4 - 13) | 1.5 (1.0 - 2.0) | -0.3 (-0.4 - -0.3) | -1.0 (-2.0 to 0.0) | 0.04 (0.00 - 0.07) |
| phloretin (n=8) | 1.1 (-0.5 - 2.1) | 11 (8 - 14) | 2.0 (1.8 - 2.2) | -0.3 (-0.4 - -0.2) | -1.0 (-1.2 to -0.8) | 0.03 (0.01 - 0.03) |
| ritonavir (n=8) | 0.9 (-0.7 - 1.6) | 8 (7 - 14) | 2.0 (1.8 - 2.0) | -0.1 (-0.5 - 0.0) | -1.0 (-2.0 to -1.0) | 0.05 (0.03 - 0.06) |
| **Kruskal-Wallis *p*-value** | 0.93 | 0.74 | 0.81 | 0.64 | 0.73 | 0.47 |

| **Treatment** | **Δ Plasma urea**  mmol L^−1^ | **Δ Mean arterial pressure**  mmHg | **Δ Heart rate**  min^−1^ | **Δ Blood hemoglobin**  g L^−1^ | **Δ Blood erythrocyte volume fraction** |
| --- | --- | --- | --- | --- | --- |
| 1.5% Glucose |  |  |  |  |  |
| Control (n=8) | 0.1 (-0.1 - 0.2) | -9 (-30 - 0) | -38 (-81 to -1) | -7 (-8 to -4) | -0.02 (-0.03 to -0.01) |
| GLUT1i (n=8) | 0.1 (-0.1 - 0.3) | -4 (-8 - 1) | -34 (-63 to 4) | -4 (-7 to -3) | -0.01 (-0.02 to -0.00) |
| phloretin (n=8) | 0.2 (-0.1 - 0.3) | -5 (-26 - 3) | -20 (-33 to -14) | -2 (-7 to 0) | -0.01 (-0.02 to 0.00) |
| ritonavir (n=8) | 0.1 (-0.2 - 0.3) | -4 (-17 - 7) | -10 (-35 to 22) | -6 (-7 to -2) | -0.02 (-0.02 to -0.01) |
| **Kruskal-Wallis *p*-value** | 0.83 | 0.60 | 0.55 | 0.65 | 0.28 |

BAY-876, selective inhibitor of GLUT1. TCO2, total carbondioxide. iCa, ionized calcium

**Supplemental Table S2.** Three-pore model parameters

| **Three-pore model parameters** | **Value** |
| --- | --- |
| Small pore radius (r_s_) (Å) | 43 |
| Large pore radius (r_L_) (Å) | 250 |
| Fractional small pore UF coefficient (α_s_) | 0.9 |
| Fractional large pore UF coefficient (α_L_) | 0.08 |
| Fractional transcellular UF coefficient (α_c_) | 0.02 |
| Unrestricted pore area over unit diffusion distance for small pores (A0/ΔX)s (cm) | 488 |
| Peritoneal lymph flow (L) (mL/min) | 0.001 |
| Transperitoneal oncotic pressure gradient (Δπ_prot_) (mmHg) | 16 |
| Fill volume (mL) | 20 |
